# Enhancer Activity-informed Gap GEne Regulatory Network (EAGER) to model *Drosophila* gap gene expression on the entire anterior-posterior (A-P) axis

**DOI:** 10.64898/2026.09.15.751584

**Authors:** Razeen Shaikh, Sebastiano Busato, Sadia Siddika Dima, Cranos Williams, Gregory T. Reeves

## Abstract

1.

Across metazoa, morphogen gradients differentially regulate gene expression and activate a spatially distinct program to specify body axis development. The *Drosophila* gap gene network, initiated by maternal morphogen Bicoid, is one of the most well-studied systems. Several regulatory interactions act synergistically to produce distinct gap gene expression patterns along the anterior-posterior (AP) axis of the blastoderm stage *Drosophila* embryo to ensure the proper segmentation of the larval and, eventually, adult stage fly. Several mathematical models have been proposed to summarize the interconnectivity of gap gene regulatory elements and predict expression in mutant systems. However, these models have not successfully predicted the gap gene expression profile over the entire AP axis. Here, we present an <u>E</u>nhancer <u>A</u>ctivity-informed <u>G</u>ap G<u>E</u>ne <u>R</u>egulatory network (EAGER) model that incorporates enhancer activity-driven differential regulation along the AP axis, which successfully summarizes gap gene expression patterns over the entire AP axis. We validated the predictions of EAGER on *Kr* mutants and performed a comprehensive parametric sensitivity and identifiability analysis to evaluate the robustness of the EAGER model fits and predictions. We also propose a reduced version of the model, rEAGER, which identifies a minimal set of regulatory interactions and successfully summarizes the gap gene interactions over the entire AP axis. Our results suggest that the expression driven by individual enhancers must be accounted for in models of developmental pattern formation.

Graphical abstract.
Enhancer Activity-informed Gap GEne Regulatory network (EAGER) to model Drosophila gap gene expression on the entire AP axis.

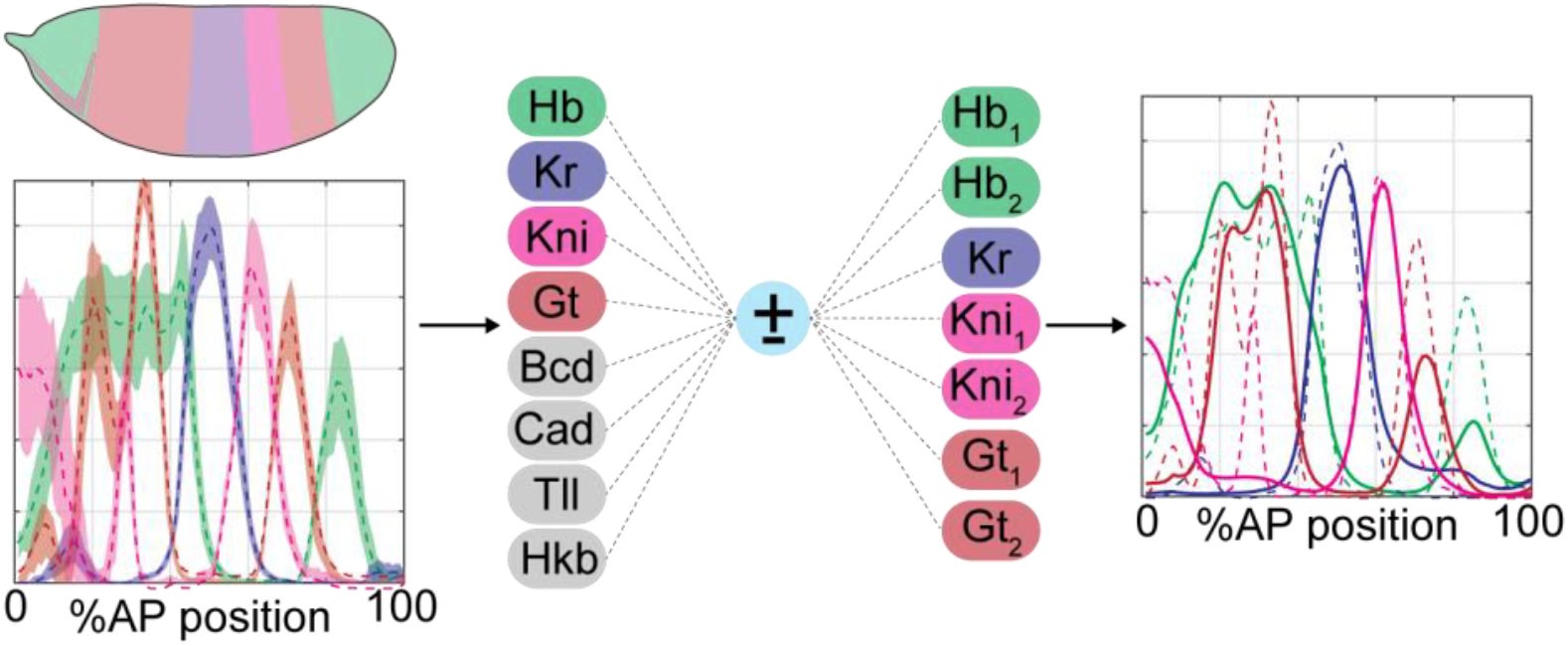

## 3. Introduction & Background

For proper metazoan development, the process of body axis patterning and segment determination is precisely regulated by a complex web of gene interactions called gene regulatory networks (GRNs). At the DNA level, this regulation is achieved by transcription factors binding to their cognate sequences, which are often clustered together into regions known as cis-regulatory elements or enhancers. At the body axis level, the initial positional information is often dictated by diffusible signaling proteins known as morphogens, which have a graded concentration profile in space, and to which cells respond in a concentration-dependent fashion^1-4^. In *Drosophila*, early patterning occurs in the syncytial blastoderm, with nuclei on the surface undergoing multiple synchronous rounds of rapid nuclear division without cytoplasmic division. At this stage, as the early embryo is considered a multinucleate single cell, intracellular transcription factors can act as morphogens by diffusing throughout the length of the blastoderm.

The process of segment determination in the early *Drosophila* embryo has been the subject of significant interest over the last three decades, with dozens of publications attempting to elucidate molecular and genetic components and their regulatory effects on downstream targets (reviewed in^5-8^). A tiered axis development program is initiated by maternally deposited coordinate gene products, including *bicoid (bcd), hunchback (hb), and nanos (nos)*, followed by zygotic gap gene activation, including *hunchback* (*hb*), *Krüppel* (*Kr*), *knirps* (*kni*), and *giant* (*gt*). The broadly expressed maternal gradients regulate overlapping zygotic gap gene domains, which initiate the segmentation process that is carried throughout the lifetime of the fly. In this context, the gap gene network plays a crucial role in generating discrete spatial patterns of gene expression from morphogen cues ^9, 10^.

The zygotic gap genes receive their initial regulatory inputs primarily through anterior, posterior, and terminal maternal systems^11^. In the anterior system, *bcd* mRNA is anchored near the anterior pole; upon translation, the Bcd protein diffuses from the anterior pole, forming an anterior-to-posterior (AP) gradient that spans most of the embryo length in an exponential-like decay^12-14^. Bcd translationally represses Caudal (Cad), establishing a posterior-to-anterior Cad gradient^15^. The posterior system includes maternal Nanos (Nos), which represses the translation of maternal *hb*, forming an anterior Hb gradient^16^. Finally, the terminal system includes Huckebein (Hkb) and Tailless (Tll), which are regulated by the Torso:mitogen-activated protein kinases (Tor:MAPK) pathway, allowing for Tll and Hkb expression at the anterior and posterior poles of the embryo^17^.

The zygotic gap gene expression produces gap proteins, Hunchback (Hb), Krüppel (Kr), Knirps (Kni), and Giant (Gt), which are transcription factors and can cross-regulate other gap proteins, thus forming the gap gene regulatory network^18^. These cross-interactions facilitate sharp domains of gene expression patterns, which stand in stark contrast to the more graded distributions of the maternal cues^19-23^. Ultimately, these sharp boundaries of spatially defined expression domains dictate the formation of patterns and segmentation in the adult fly.

The advent of quantitative biology tools and advanced imaging techniques has catalyzed a thorough investigation of the spatiotemporal dynamics of gap gene proteins. Several quantitative studies employing techniques including live imaging, fluorescence recovery after photobleaching, fluorescence correlation spectroscopy, fluorescence lifetime analysis, and optogenetics have been reported for Bcd alone^24-32^. In turn, these quantitative data generated enthusiasm to develop mathematical models that summarize the gap gene expression profiles and predict them in mutant systems^33-46^.

Early mathematical models of the gap gene network assumed that proteins in the maternal and terminal systems are static, which is incorrect, as these inputs are dynamic^40^. These models were refined by treating the maternal and terminal systems only as inputs and including terminal *hkb* expression^47^. These models also assumed a stationary Bcd expression profile; however, the maternal inputs, including Bcd and Cad, are dynamic, on the same timescale as the four gap genes^40-42, 48^. In addition to these biological discoveries, parameter estimation tools, including improved evolutionary algorithms, have been developed to efficiently solve the inverse problem of fitting the model to the data^49-51^. Despite these conceptual advances, most modeling efforts, including ours^51^, have been unable to properly recapitulate gap gene expression patterns over the entire AP axis, and thus limited the spatial domain to a subset (35-92%^42^) of the AP axis. This truncation of the AP axis remains perhaps the biggest conceptual lacuna in modeling the gap gene network, and may indicate that a single regulatory ruleset for each gap gene is not sufficient.

Recent developments suggest that each gap gene is regulated by multiple enhancers, and their combinatorial effect, also referred to as “enhancer synergy”, ensures precise spatial patterning, and each enhancer alone cannot reproduce precise gap expression patterns^52-55^. It has been shown that, while the multiple enhancers for a gene often have partially redundant function, the loss of one of a gene’s enhancers leads to perturbed patterns of gene expression, suggesting that they work synergistically to ensure precise spatial limits of gene expression ^55^. Additionally, Kvon et al. provided a genome-wide spatial roadmap to identify *in vivo* enhancer activity of over 7,700 *Drosophila* enhancers in developing embryos^56^. Previous attempts to account for the presence of multiple *gt* enhancers in the gap gene expression model have focused their efforts on either incorporating the enhancers for one gap gene only (*gt*) or limiting the spatial domain along the A-P axis^44,57^. Despite their successes in modeling and predicting gap gene expression, several limitations include reduced model complexity and flexibility of gap gene interactions, potentially highlighting the need for improved models^44, 57^. However, no prior efforts have accounted for differential gap gene regulation through the activity of multiple enhancers of each gene. The scope and resolution of recent findings provide a basis for an upgrade to the classical modeling approach and the need to account for the activity of multiple enhancers as an additional component to the existing regulatory framework.

In this manuscript, we propose a modeling framework, <u>E</u>nhancer <u>A</u>ctivity-informed <u>G</u>ap G<u>E</u>ne <u>R</u>egulatory network (EAGER), which decouples the transcriptional output of experimentally reported gap gene enhancers and allows for each gap gene to be differentially regulated by multiple rule sets of gap protein interactions along the AP axis. The EAGER framework allows for spatial flexibility in modeling gap gene interactions. We circumvent complications associated with the increase in the number of parameters by implementing a computationally efficient evolutionary algorithm to solve the equations, and by employing regularization strategies to reduce the number of parameters post hoc. We show that our model correctly reproduces gap gene patterns on the entire AP axis for nuclear cycles 13 and 14; further, the resulting parameters are in line with expected interaction strengths, given prior experimental results and *in silico* sequence analysis. Finally, our results correctly reproduce gap gene expression in *Kr* mutants, highlighting the predictive power of our approach.

## 4. Results

### 4.1. The <u>E</u>nhancer <u>A</u>ctivity-informed <u>G</u>ap G<u>E</u>ne <u>R</u>egulatory (EAGER) network summarizes gap gene expression over the entire AP axis

The genes composing the Gap gene network are each regulated by multiple enhancers, and the synergistic activity of these multiple enhancers contributes to the final gene expression pattern ^54,58^. However, previous models of the Gap gene network neglected the presence of multiple enhancers for each gene. Therefore, to model the Gap gene network, we propose an <u>E</u>nhancer <u>A</u>ctivity-informed <u>G</u>ap G<u>E</u>ne <u>R</u>egulatory (EAGER) network, which incorporates the activity of multiple enhancers for each gene along the anterior-posterior (AP) axis. The distinct activity of these enhancers, including differential spatial domains, has been visualized in blastoderm-stage embryos in which gene expression is driven by individual gap gene enhancers ^52, 53, 59^. We used a database^14, 47^ of these images to inform our decision to model the spatial domains of gap gene expression driven by their distinct enhancers during nc 13 and nc 14 (see Methods). The database images and other literature suggest that, for each of *hb, gt*, and *kni*, at least two enhancers drive spatially non-overlapping expression patterns, whereas for the fourth gap gene, *Kr*, the same spatial expression pattern is driven by both enhancers ^52-55, 59, 60^. We integrated these data into the mathematical model by allowing distinct enhancers, which are active in spatially distinct regions of the A-P axis, to act independently in decoding the system inputs and regulating the activity of other gap genes to produce patterns of gene expression over the entire A-P axis (Figure **1**). The model also incorporates the dynamics of maternal inputs of Bcd and Cad, and terminal systems of Tll and Hkb ^40-42, 47^.

**Figure 1.**
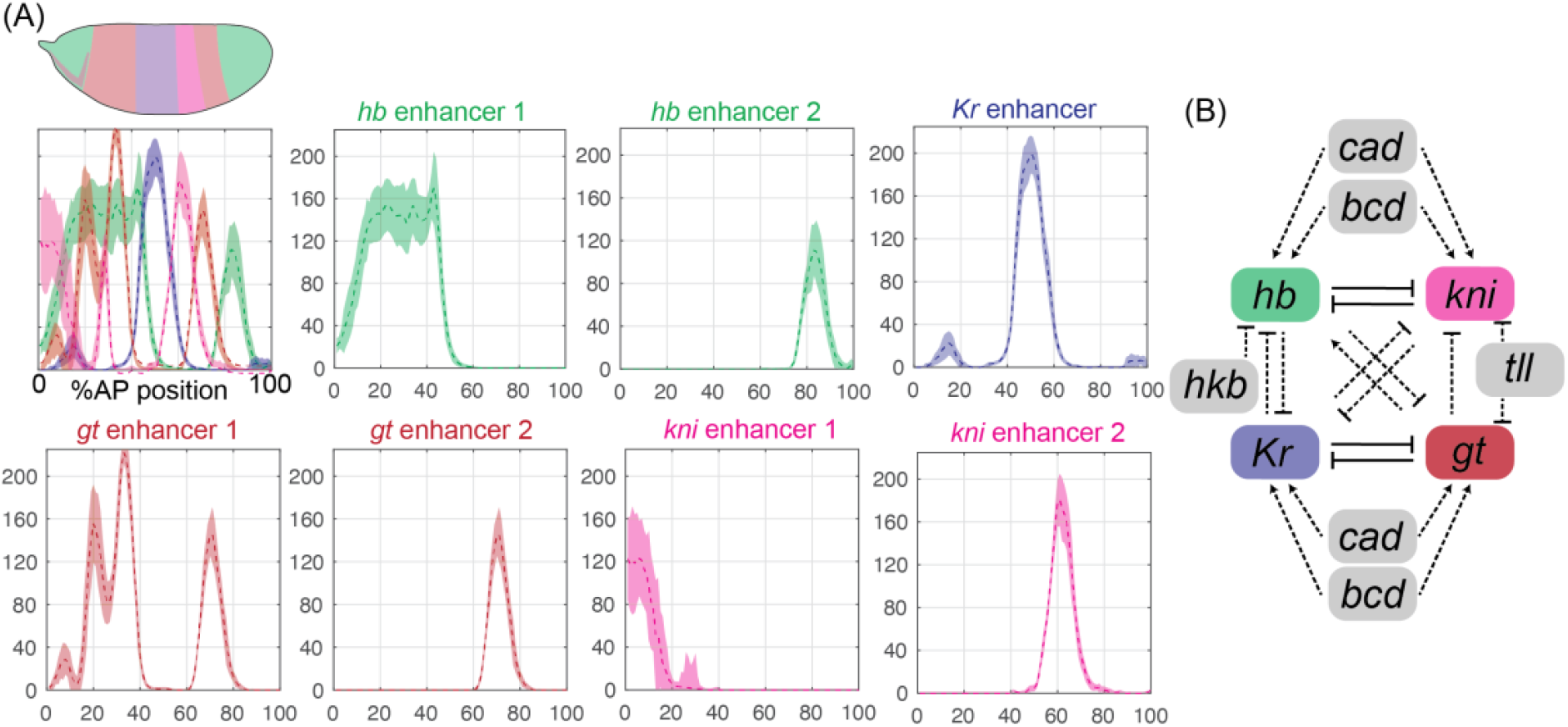
Enhancer driven gap gene expression during nc14 over the entire AP axis. **(A)** Gene expression profiles for the four gap genes over the entire AP axis. In the Stark Lab dataset, the expression of *hb, gt*, and *kni* is driven by two enhancers, whereas *Kr* is driven by one enhancer. **(B)** The canonical gap gene network in *Drosophila* blastoderm embryo.

We fit this model to gap gene expression profiles, using an evolutionary optimization algorithm, ISRES+, and obtained ∼40 parameter sets with acceptable goodness of fit (GOF) values^51, 61^. The simulated gene expression profiles (black curves) with the best-fit model parameters are consistent with the experimental expression profile (gray shaded region) for the anterior Hb profile, Kr peak, the two Gt peaks (Figure **2A-C**) in nc 13; and the two Hb peaks, prominent Kr peak, three prominent Gt peaks, the anterior Kni expression (Figure **2H-K**). While the model outperforms previous models in correctly delineating the anterior and posterior boundaries of several gap protein expressions, it fails to reproduce less prominent Kr and Gt peaks and the middle Kni peak (Figure **2I-K**). The model also fails to correctly delineate the trough between the two anterior Gt peaks, perhaps due to its inability to identify the Kni peak in that location (Figure **2J-K**). It has been previously noted that the discrepancies in the total expression level are less important since these data are arbitrary due to relative protein concentrations^62^.

**Figure 2.**
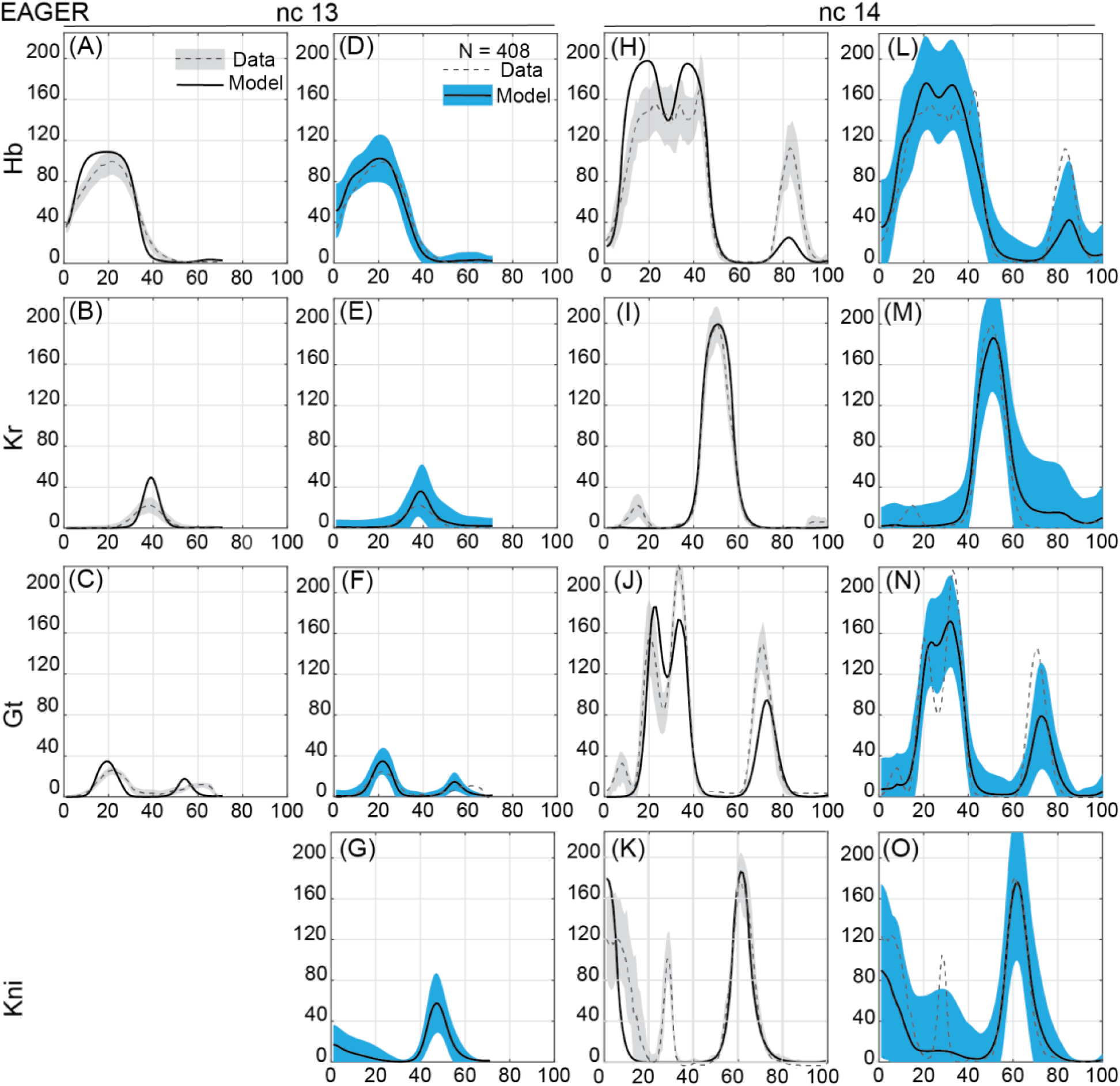
Gap gene expression predicted over the entire AP axis. **(A-C)** The predicted expression profiles of Hb (A), Gt (B) and Kr (C) during nc 13 with the best fit parameter set and the experimental data (average dashed line, standard deviation gray), **(D-G)** The average expression profile (black) and the confidence interval (blue) of the predicted expression profiles of Hb (D), Gt (E), Kr (F) and Kni (G) during nc 13, **(H-K)** The predicted expression profiles of Hb (H), Gt (I), Kr (J) and Kni (K) during nc 14 with the best fit parameter set and the experimental data (average dashed line, standard deviation gray), **(L-O)** The average expression profile (black) confidence interval (blue) of the predicted expression profiles of Hb (L), Gt (M), Kr (N) and Kni (O) during nc 14.

By previously noted definitions, the transcriptional weights (interaction parameters, T) are ill-determined if their estimated values are not strictly greater than or less than zero, and span over the entire (-1,1) range, meaning that the model is unable to identify the regulatory action of one gene on another. Our results indicate that most of the estimated parameters are not ill-determined (Supplementary Section **2.1**)^62^. Despite the improved performance of EAGER, the model contains 68 variable parameters. We needed to verify that the gap expression is robust to parameter perturbations, unlike previous work, where the posterior Gt peak has been reported to be sensitive to perturbations ^43^. To evaluate the robustness of these results and to account for the sloppiness in systems biology models, we generated 408 “shuffled” combinations of the 6 parameter sets that fit the data nearly well (referred to as “well-fit” parameter sets) and simulated the model output at all these combinations ^61, 63, 64^. The average simulated expression profile from the 408 simulations (black curve) and its confidence interval (blue shaded region) for each of the gap genes in nc 13 (Figure **2D-G**) and nc 14 (Figure **2L-O**) reveal that the EAGER network robustly summarizes gap gene expression patterns during nc13 and nc14 within one standard deviation of the experimental data (Figure **2**). Not only does the model consistently identify all prominent expression patterns but it also begins to identify the less prominent Kr and Gt (Figure **2M-N**) anterior peaks, and the middle Kni peak (Figure **2O**).

In summary, using the “best-fit” parameter sets, the model can correctly generate nearly all gap expression profiles in nc13 (Figure **2A-C**) and nc14 (Figure **2H-K**); and, by generating 408 combinations of the 6 well-fit parameter sets, we show that the model can robustly identify prominent expression patterns and can identify the less prominent peaks as well. Most importantly, the EAGER network model is the first model to summarize gap gene expression profiles over the entire AP axis.

### 4.2. The EAGER network can successfully predict the gap gene expression in *Kr* mutants

Most models of gap gene expression fail to predict the expression pattern in mutant backgrounds ^43, 47, 57^. To validate the predictions of the EAGER network, we used the estimated parameters to predict gap protein expression profiles in the *Kr* null mutant background during time class 3 (tc 3) and time class 7 (tc 7) of nc 14 ^37^.

Proper expression of zygotic *Kr* is essential for embryogenesis to the extent that alterations in *Kr* activity affect the segmentation pattern and cause lethality^65^. The *Kr* expression pattern is transcriptionally and post-transcriptionally regulated by nearly all the other gap gene products^66-68^. Mutations in *Kr* manifest as a substantial shift in the Gt posterior domain^69^. Moreover, the auto-regulatory activity of *Kr* and its cooperativity with *hb* ensure sharp pair-rule gene expression bands and altered *Kr* dosage affects the spatiotemporal dynamics of *eve* stripe formation^68, 70-73^. The interactions among gap proteins, including *Kr* and *gt*, which are expressed in complementary domains, are mutually repressive and strong^74^. In contrast, the interactions among gap proteins, including *Kr* and *hb*, which are expressed in overlapping domains, are relatively weak^75^. For these reasons, validating the predictions of EAGER in *Kr* mutants is crucial.

EAGER predictions with the best-fit parameter set normalized to the maximum intensity of gap protein expression profiles for Hb, Gt, and Kni indicate that the EAGER model identifies almost all of the gene expression peaks, including correct placement of anterior and posterior boundary locations in tc 3 and tc 7 of *Kr* mutants (Figure **3A-C**, **3G-I**). However, the best-fit EAGER predictions do not perfectly delineate the anterior boundary of posterior Gt (Figure **3B**), the posterior boundary of anterior Kni in tc 3 (Figure **3C**); and the posterior boundary of posterior Hb (Figure **3G**), the posterior boundary of anterior Gt (Figure **3H**), and the boundaries of the posterior Kni (Figure **3I**). These predictions are on par with previous attempts to model gap gene expression in *Kr* mutants^57^. Surprisingly, the average gap expression profile with the 408 “shuffled” combinations of parameters fares better than the best-fit prediction alone (Figure **3D-F, 3J-L**). There are clear improvements in Hb and Gt profile predictions in both tc 3 and tc 7, including peak heights. Despite the model failing to cleanly predict Kni expression, the high variability in the anterior Kni predictions captured by the confidence interval hints that, under some parameter sets, the EAGER model at least partially captures the Kni peak. We expect that further optimization of the EAGER network parameters could facilitate precise and accurate Kni prediction.

**Figure 3.**
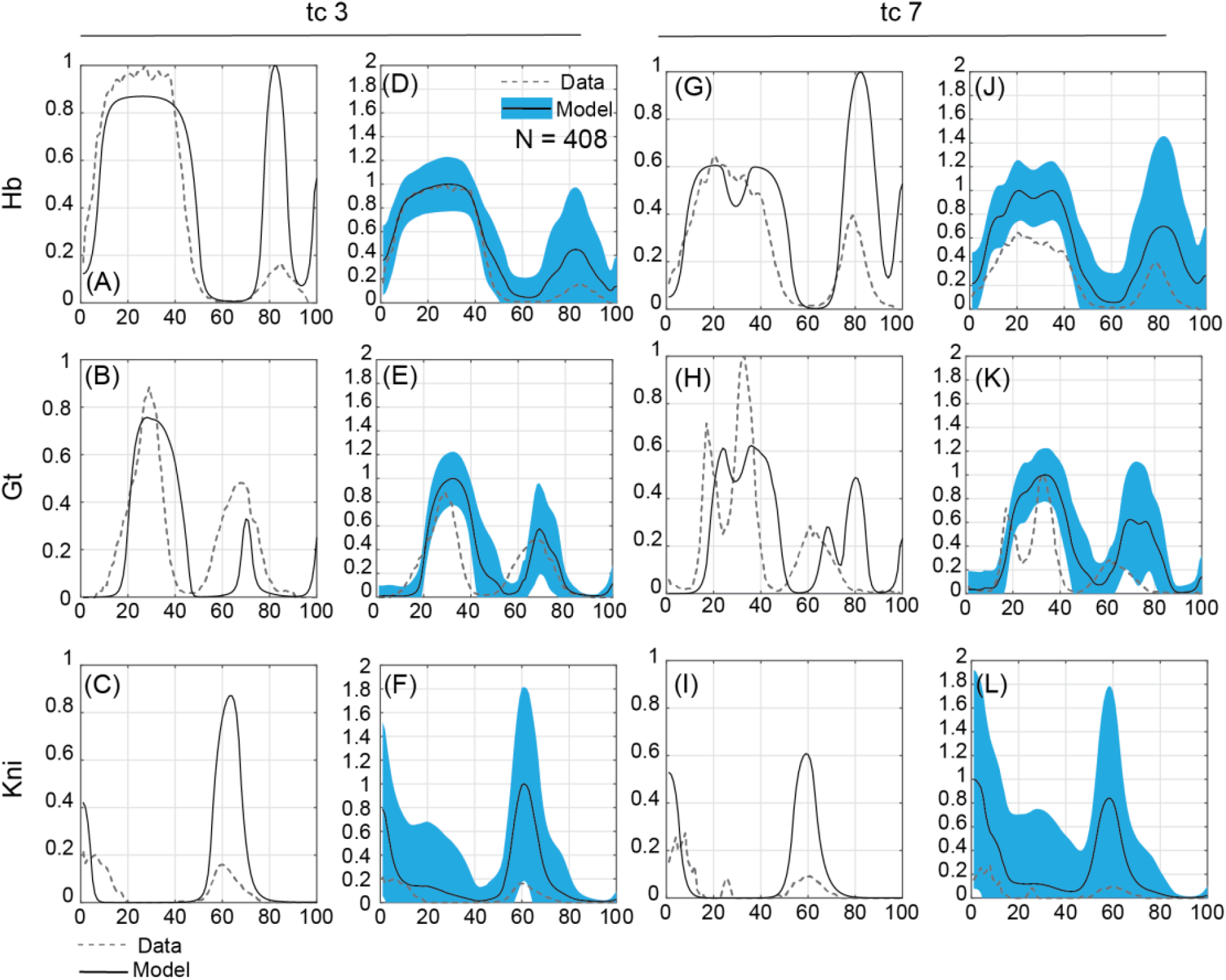
Model Validation on gap gene expression in Kr mutants. **(A-C)** The tc 3 gap gene expression in Kr null background predicted by the best-fit parameter set of the EAGER model for Hb (A), Gt (B) and Kni (C), **(D-F)** The average expression profile (black) and confidence interval (blue) of the gap gene expression predicted by the EAGER model for Hb (D), Gt (E), and Kni (F), **(G-I)** The tc 7 gap gene expression in Kr null background predicted by the best-fit parameter set of the EAGER model for Hb (G), Gt (H) and Kni (I), **(J-L)** The average expression profile (black) and confidence interval (blue) of the gap gene expression predicted by the EAGER model for Hb (J), Gt (K), and Kni (L)

### 4.3. The reduced-EAGER (rEAGER) model summarizes gap gene expression profiles

While the EAGER network model outperforms other gap gene models in the literature, it incorporates a large number of variable parameters (68). Models in systems biology are “sloppy,” such that many parameter sets fit the data equally well, resulting in varied predictions and impeding our ability to harness the predictive power of the model^61, 63, 64^. To reduce the number of model parameters, we implemented two regularization strategies: Ridge regression (Equation **4**) and LASSO (least absolute shrinkage and selection operator; Equation **3**)^76^. These regularized linear regression variants attempt to simultaneously minimize the GOF and reduce the number of variable parameters (Supplementary Section **3**).

We first implemented the LASSO regularization strategy by adding the norm of the transcriptional weights to the GOF function, which, in addition to the difference between the model fit and experimental data, also minimized the transcriptional weights (Equation **3**). We ran the evolutionary algorithm, ISRES+, at least 100 times with the LASSO-variant of the EAGER GOF function “EAGER-LASSO”, of which 21 results converged on a solution with a GOF value between 1.9 and 4.2. As such, applying LASSO regularization indeed minimized the transcriptional weights (Supplementary Section **2.3**) without significantly compromising EAGER’s ability to summarize gap gene expression over the entire AP axis (Supplementary Section **3.1**). Using EAGER-LASSO did not compromise the nc 13 fits. However, with EAGER-LASSO, we lost the ability to identify the posterior Hb peak, the anterior Gt trough, and the middle Kni peak as prominently as EAGER. This shows that we can summarize gap gene expression without using 68 variable parameters, and perhaps several of these interactions are nonexistent.

As an alternative, we implemented the Ridge regularization strategy by adding the square of the norm of the transcriptional weights to the EAGER GOF function, “EAGER-Ridge” (Equation **4**). The choice of the GOF function plays a significant role in performing efficient parameter estimation, so it was necessary to test at least two variants^45, 77^. Once again, we ran ISRES+ at least 100 times, of which 35 instances resulted in an EAGER-Ridge GOF value between 1.6 and 4.1. Relative to LASSO, Ridge is less aggressive in forcing the transcriptional weights to a trivial value (Supplementary Section **2.3**). While EAGER-Ridge is worse than EAGER-LASSO at identifying Kr in nc 13, it retains the posterior Hb peak (Supplementary Section **3.2**).

After finding that LASSO is superior to Ridge at balancing the trade-off between minimal transcriptional weights and maximizing the ability to generate good fits, we chose to perform further analysis on EAGER-LASSO. Our goal was not just to minimize the transcriptional weights but to completely remove them and reduce the number of model parameters from a staggering 68. To do this, we set to zero all transcriptional weights less than an absolute cut-off of 0.05, resulting in the removal of 15 transcriptional interactions. We also tested other values of the cut-off, which removed more connections without significantly improving goodness of fit. After removing these 15 weak interactions to create a reduced-EAGER (rEAGER) model (Figure **4D**), we found that the best-fit does not specify the peak characteristics as accurately as EAGER, including the Kr and Gt nc 13 profiles, the posterior Hb peak, and the shape of Kr in nc 14. Interestingly, rEAGER gains an ability to specify the anterior Kni even better. However, the confidence intervals for rEAGER and EAGER are either overlapping nearly perfectly (Hb and Kni) or further constrained (Kr and Gt). Thus, by removing 15 parameters, we essentially have the same number of parameters as all the other gap gene models and yet retain EAGER’s ability to model gap gene expression over the entire AP axis.

**Figure 4.**
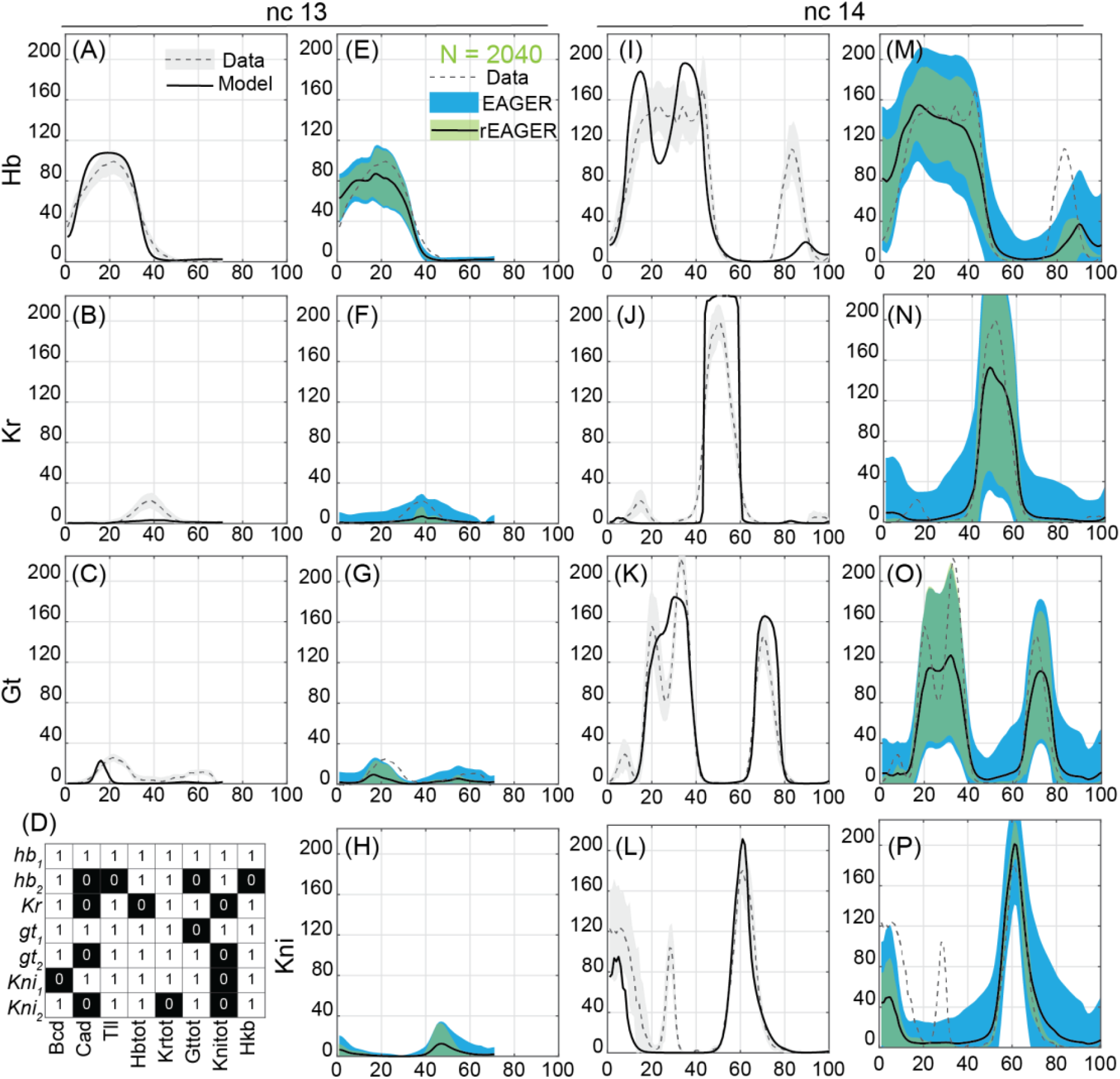
Gap gene expression predicted by rEAGER model over the entire AP axis. **(A-D)** The predicted expression profiles of Hb (A), Gt (B) and Kr (C) during nc 13 with the best fit parameter set and the corresponding heatmap of removed interactions (D), **(E-H)** The average expression profile (black) and the confidence interval of the predicted expression profiles of Hb (E), Gt (F), Kr (G) and Kni (H) during nc 13, **(I-L)** The predicted expression profiles of Hb (I), Gt (J), Kr (K) and Kni (L) during nc 14 with the best fit parameter set, **(M-P)** The average expression profile (black) and the confidence interval (green) of the predicted expression profiles of Hb (M), Gt (N), Kr (O) and Kni (P) during nc 14.

The maternal proteins Bcd and Cad are largely known as activators of all gap genes^44^. Still, rEAGER predicts that Bcd does not regulate anterior Kni, and Cad does not regulate several peaks, including posterior Hb, Kr, posterior Gt, and posterior Kni. Similarly, the terminal genes Tll and Hkb are generally known to be repressors^44^, but rEAGER predicts that they do not affect the posterior Hb domain, which has been verified experimentally^47^. Other known interactions, including Cad activation of posterior Gt, are missed by rEAGER^44^. However, several known interactions, including Bcd activation of anterior and posterior Gt, Cad activation of anterior Gt, Hb repression of anterior Gt, Tll and Hkb repression of Gt, and strong Gt-Kr mutual repression, were retained by rEAGER^44^. All the gap genes except Kni are known to autoregulate their expression, and rEAGER predicted that Kni does not autoregulate itself^18^. In summary, while our data-driven analysis of the gap gene network may miss some regulatory connections, it nevertheless summarizes the gap profiles over the entire AP axis and also predicts the existence of several known gap gene interactions correctly.

### 4.4. The rEAGER network can successfully predict the gap gene expression in *Kr* mutants

We validated the predictions with EAGER-LASSO and EAGER-Ridge on *Kr* null mutants, and despite EAGER-Ridge’s ability to fit the data better than EAGER-LASSO, its ability to predict gap expression in *Kr* mutants is relatively inferior (Supplementary Section **3.3**). The best-fit results for both the regularization strategies look similar; the confidence intervals for the Ridge for posterior Hb and LASSO for posterior Kni have a large variability, which we deem a disadvantage in making robust predictions.

We validated the predictions of the rEAGER network for *Kr* null mutants with the best-fit parameter sets described in the previous section (see Methods). The rEAGER best-fits and average expression profiles with confidence intervals are in fair agreement with the *Kr*-mutant data, such that all the peak locations overlap with the normalized expression data for Kr mutants (Figure **5**). In addition, we are finally able to predict the second Kni peak, which so far has been excluded by our fits (Figure **5L**).

**Figure 5.**
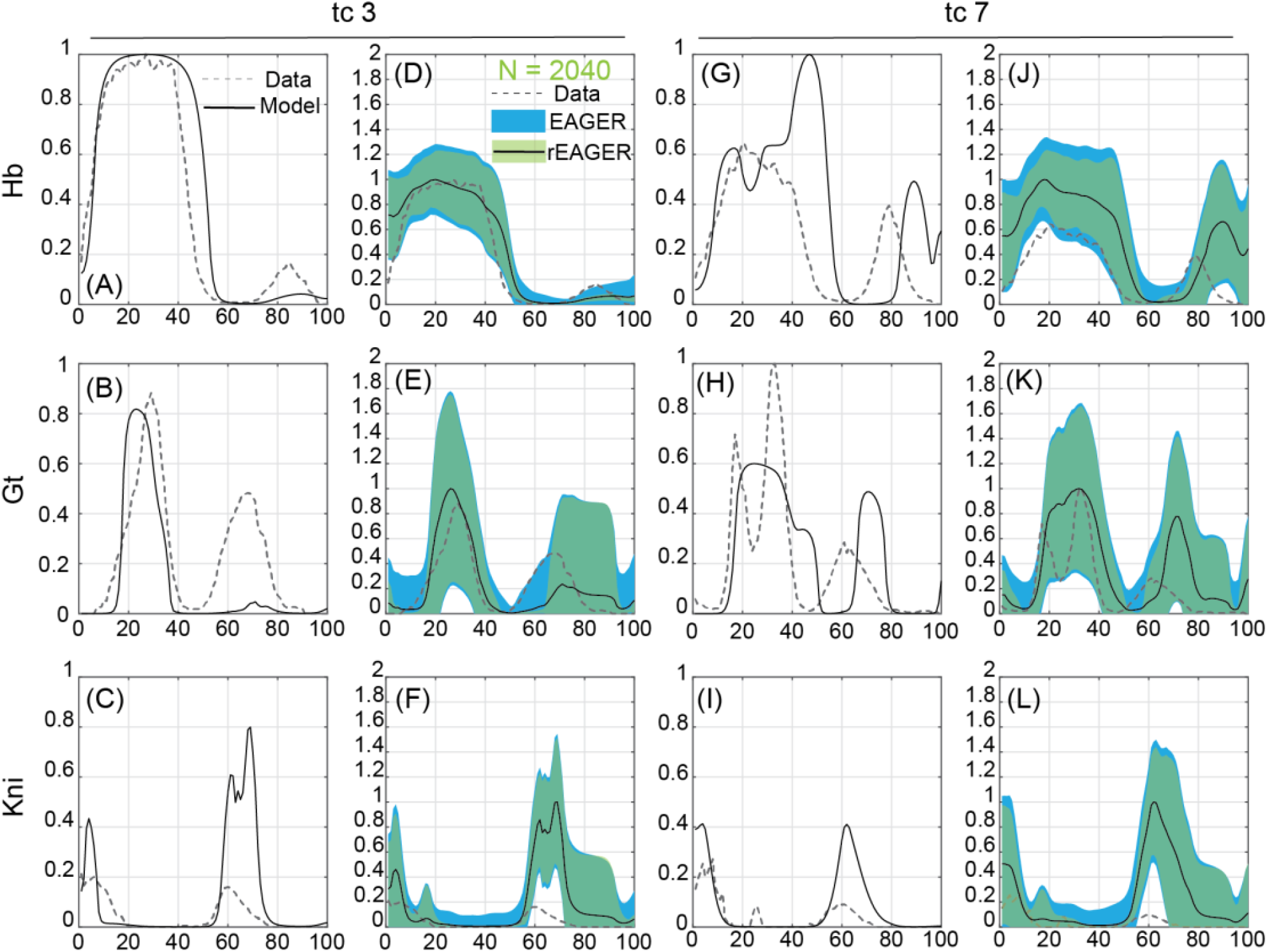
Gap gene expression predicted by rEAGER model over the entire AP axis. **(A-C)** The predicted expression profiles of Hb (A), Gt (B) and Kr (C) during nc 13 with the best fit parameter set, **(D-G)** The average expression profile (black) and the confidence interval (green) of the predicted expression profiles of Hb (D), Gt (E), Kr (F) and Kni (G) during nc 13, **(H-K)** The predicted expression profiles of Hb (H), Gt (I), Kr (J) and Kni (K) during nc 14 with the best fit parameter set, **(L-O)** The average expression profile (black) and the confidence interval (green) of the predicted expression profiles of Hb (L), Gt (M), Kr (N) and Kni (O) during nc 14.

The parameter reduction via regularization, going from EAGER to rEAGER, partially confirms the reliability of our model. The interactions retained point to several well-established interactions^44^, including mutual repression between *gt* and *Kr*, repression of anterior *gt* by Hb, Bcd activation of anterior and posterior *gt*, Cad activation of anterior *gt*, terminal repression of *gt* by Tll and Hkb, and autorepression of all gap genes (except *kni*, as expected^18^). In addition, several of the departures from the canonical network interactions highlighted by the parameters pruned by rEAGER are in fact consistent with recent experimental work. rEAGER predicts that Bcd does not activate anterior *kni*, in contrast with the accepted view that Bcd and Cad are general gap gene activators^44^. However, a recent study utilizing optogenetic control of Bcd in live embryos has identified a non-canonical suppressive relationship of Bcd on kni, with kni transcription initiating rapidly after Bcd depletion^32^. Similarly, rEAGER indicated no effect of Cad on several posterior domains, including posterior Hb, Kr, and Gt posterior peaks. This is in alignment with the work of Verd and colleagues^40, 78^, who showed that the shift rate of posterior gap domains is largely independent of maternal Cad levels, indicating a weaker effect of Cad on that region of the A-P axis.

### 4.5. Parameter Sensitivity Analysis

In the previous sections, we have confirmed that the model-predicted gap gene expression profiles are not sensitive to parameter combinations and that our estimated parameter ranges are robust. To further evaluate the robustness and validity of our model predictions with the best fit parameter set (Figure **2H-K**), we performed a parametric sensitivity analysis by locally perturbing each model parameter (*δ*∼1%) and calculating the percentage change in peak height (H), peak location (L), and anterior (A) and posterior (P) boundary locations. The goal is to determine the extent to which individual parameters influence peak characteristics and identify non-canonical gene interactions that contribute to shaping gap gene expression (Supplementary Section **4**).

First, if small local perturbations in parameters have a strong effect on peak characteristics, then our results are likely sensitive to parameter combinations, and the predictive power of the model would be low. The analyses indicate that local perturbations have a minor effect (within ∼2%) in defining all the Kr and Kni peak characteristics. The local perturbations in parameters have a minor effect on nearly all the Gt and Hb peak characteristics, except in a few instances. Our previous analysis (Figure **2L**,**M**) and this parameter sensitivity analysis highlight the limitations of our ability to predict the height of the second Hb peak and delineate the boundaries of the anterior Gt peaks, respectively.

Second, with this local analysis, we can test whether perturbing the interactions between gene_a_ and gene_b_ affects the expression of gene_c_. Our analysis predicts several regulatory interactions that indirectly affect peak characteristics. For example, we found the posterior Kr boundary is sensitive to the effect of Cad regulation of the second *kni* peak, the Gt peak height and location are sensitive to the strength of several regulatory inputs to the first *hb* domain, and the Kni posterior profile is sensitive to Cad regulation of *Kr*.

Our work exemplifies the utility of data-driven analysis to model gap gene expression on the entire AP axis, predict the gap expression in mutants, identify key regulatory interactions, and propose the effect of perturbing gap interactions locally on the complete expression patterns.

## 5. Discussion

Over the last two decades, *Drosophila* embryonic development has been the subject of extensive quantitative analysis. Improved high-throughput genetic perturbation, imaging, and analysis platforms have enabled the generation of large quantitative datasets^56, 79, 80^. However, translating these quantitative datasets into predictive models has been challenging. Numerous advances in simulation techniques and integration of biological modeling constraints have improved the ability to summarize gap gene interactions and predict gap gene expression patterns^40, 41, 44, 45, 47, 48^. However, to the best of our knowledge, no effort has mathematically reproduced gap gene expression patterns over the entire A-P axis. In this paper, we report EAGER: an enhancer activity-informed gap gene regulatory network, which summarizes gap gene expression over the entire A-P axis and can successfully predict the gap gene expression in a mutant background.

Transgenesis and quantitative imaging show that multiple enhancers regulate each gap gene, and the gap gene pattern is a combinatorial outcome of synergistic enhancer activity. EAGER utilizes a genome-wide spatial mapping of *Drosophila* enhancers and their activity to inform the classical synthesis-diffusion-degradation formulation of the gap gene regulatory network. This enabled us to deconvolve the role of individual enhancers in driving gap gene expression in distinct spatial locations, such that two distinct enhancers can additively drive Hb, Gt, and Kni patterns. We fit EAGER to the gap gene expression data in nc13 and nc14 using an evolutionary strategy, ISRES+^51^, and obtained several parameter sets with high goodness-of-fit that reproduced the gap gene expression pattern with high fidelity and narrow confidence intervals. Additionally, we validated the predictions of EAGER against tc3 and tc7 gap patterns in *Kr* mutants, and despite a few inaccuracies, EAGER outperforms any other model.

With EAGER, the core of our approach rests on two deliberate modeling decisions. The first is that expression patterns of Hb, Gt, and Kni can be separated into spatially resolved, enhancer-driven components, according to the portion of the A-P axis in which each enhancer is predominantly active. This framing is in partial contrast to the prevailing opinion that multiple enhancers are substantially biologically redundant, driving overlapping expression patterns, ensuring biological robustness^81^. We do not reject that such overlaps exist; however, our approach is substantiated by the observation that *Drosophila* enhancers can express genuinely distinct spatiotemporal patterns both in the gap gene network^44, 60^ as well as in other gene regulatory networks^82^.

Our second deliberate decision is to treat enhancer-driven expression profiles as fully additive. We choose this additivity as a simplification of biological reality: although earlier work treats enhancer activity as additive, more recent quantitative studies show that developmental enhancers may depart from additivity in certain situations^83^. For example, hb enhancers were found to combine additively when Bicoid is low, but sub-additively when Bicoid is high^54^. Further, a recent study of millions of enhancer pairs in S2 *Drosophila* cells revealed that developmental gene enhancers typically act synergistically^84^. Critically, departures from additivity have been shown to concentrate precisely where enhancer activity overlaps, or where gene expression domains transition: at the boundary between the Kni and Kr domains, for example, enhancers repress one another to sharpen the expression pattern^60^. This is consistent with the areas where EAGER is least accurate: the middle Kni peak, the height of the second Hb peak, and the trough between the anterior Gt peaks fall at or near the boundaries between the enhancer-driven domains we defined, and thus, where the additivity assumption is more likely to fail. Therefore, we suggest that the inaccuracy of our model in these regions is not solely tied to parameter optimization, but rather points to enhancer dynamics not captured by our simplified assumption.

Classical interpretations and recent quantitative analysis suggest that several interactions in the gap gene network are either unidirectional or absent. However, EAGER is blind to such information. To reduce model complexity and identify key gap gene interactions, we introduced a regularization-reduced framework: rEAGER. rEAGER reproduces the gap patterns with minimal interactions and successfully predicts gap patterns in *Kr* mutant data. However, our framework failed both to identify known key interactions and to deemphasize interactions known to be insignificant. Direct quantitative measurements of endogenous and perturbed transcriptional programs would help further constrain parameters and improve predictions^32, 85^.

Models in systems biology are sloppy: large ODE models, like EAGER, are fit to sparse data, increasing parameter uncertainty and impeding predictive capability^61, 63^. We built the EAGER framework around global parameter estimation, robust model validation, confidence evaluation, model reduction through regularization, and local sensitivity analysis to enable high predictability. Despite the effort, several avenues of future research could improve the model’s predictive capabilities, including determining the role of non-canonical gap gene enhancers, cooperativity between enhancers, and the seemingly redundant role of some enhancers. Even so, without this information, the model is built on reasonable assumptions and is able to predict a wide range of wildtype and mutant gene expression patterns.

## 6. Methods

### 6.1. Gene expression and enhancer activity data

Relative protein expression data for the transcription factors Bcd, Cad, Tll, Hkb, Hb, Gt, Kni, and Kr were obtained from the FlyEx database^86^. These data are reported for nc 13 and tc 1-8 of nc 14, distributed about 6.5-8 minutes apart, with tc 8 corresponding to around 67 minutes after the 13^th^ nuclear division. This is summarized as tabular data from 125 distinct sets of images for nc 13, and 787 sets from tc 1-8 in nc14. Additionally, we utilized background-corrected data, normalized by Fast Redundant Dyadic Wavelet Transform (FRDWT)^14^, to extract the variability in expression patterns for the four gap genes. We averaged the expression of all normalized embryos to extract the canonical pattern and the standard deviation at each point on the AP axis. (Please note that at the time of publication, access to the FlyEx database appears to have been deactivated, or the servers are no longer maintained. Therefore, we have provided raw data from the database as Supplementary Table **1**.) Expression patterns for the terminal gap gene *hkb* were included as external input of the system and were obtained from prior work by Ashyraliyev *et al*.^47^

Enhancer activity data were obtained from the Stark Lab Fly Enhancers database (https://enhancers.starklab.org/)^56^ and Perry et al., 2011 (Supplementary Section 1; Figure S1A)^55^. We extracted images of all embryos with enhancer activity in the canonical gap protein expression regions of the AP axis. We filtered the resulting images to those active in developmental stages 4-6 (broadly corresponding to nc13-14) and with the “add annotation” functionality to filter the search results with terms including “gap”, “AP stripe”, “anterior”, or “posterior”. We visually examined the resulting images and oriented them by identifying the anterior and posterior poles of the early embryo.

We used MATLAB to quantify the expression patterns along the AP axis^87^. Briefly, the images were subjected to the following processes: background subtraction, identification of the location of the embryo in the image, identification of the embryo borders, image segmentation, identification of nuclei along the periphery, and finally quantification of pixel intensities in each nucleus. The resulting spatially resolved profiles of enhancer activity were compared to the canonical expression pattern of the corresponding gap gene. Areas of overlap between the expression of the gap gene and the corresponding enhancers were used to “split” the expression of the gap gene into its putative enhancer-driven components (Figure **1**). In total, two enhancers were identified for Hb, Gt, and Kni, and one for Kr.

The decision to model each gene by these specific enhancer sets was justified by prior experimental investigation of the gap genes. For Hb, a proximal and a distal shadow enhancer drive spatially resolvable activities, with the proximal being largely active in the anterior pole^55^. For gt, a prior study showed that anterior and posterior expression are driven by distinct enhancers, with one of them being responsible for a large portion of the posterior peak (between A-P 0.6 and 0.8), and the other driving expression of both peaks^52^. Similarly, kni is regulated by a proximal and a distal/shadow enhancer, with its proximal enhancer defined almost entirely within its second peak^52, 55^. In contrast, although two putative enhancers have been identified for Kr, they both drive essentially the same central domain, and their activity appears to be redundant^55^; we therefore modeled Kr through a single equation.

### 6.2. The Enhancer Activity-informed Gap GEne Regulatory network (EAGER)

#### (1) Model Architecture

To account for the influence of the two enhancers of Hb, Kni, and Gt (hereafter, subscripts “1” and “2” denote corresponding enhancer-driven expression profiles) and one Kr enhancer in specifying gap gene expression along the entire AP axis, we developed a spatiotemporal model of the intracellular dynamics of the four gap genes. We discretized in space by the 71 nuclei in nc 13 and 100 in nc 14, resulting in 71 × 7 and 100 × 7 coupled ordinary differential equations (ODEs) in nc 13 and 14, respectively. Note that the model tracks the dynamics of four gap genes but has seven species: *hb*_*1*_, *hb*_*2*_, *Kr, gt*_*1*_, *gt*_*2*_, *kni*_*1*_, *and kni*_*2*_.

Mathematically, this is formulated as,

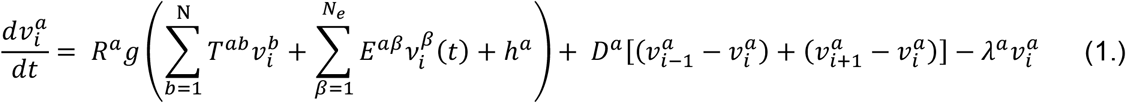

Here, *a* ≡ (*hb*_1_, *hb*_2_, *Kr, gt*_1_, *gt*_2_, *kni*_1_, *kni*_2_) denotes the model species such that the expression profile of each of the four gap genes, *b* ∈ (1, …, *N*), driven by two corresponding enhancers is additive, for instance, Hb_tot_ = Hb_1_ + Hb_2_ and so on. The three terms of the right represent protein synthesis, Fickian diffusion, and first-order protein degradation, respectively. The first term simulates protein synthesis as an activation function *g*(*u*) multiplied by the maximum synthesis rate *R*. The activation equation is a function of gap gene interactions, the spatiotemporal maternal Bcd input, the three time-varying inputs of Cad, Tll, and Hkb *β* ∈ (1, …, *N*_*e*_), and the effect of the ubiquitous transcription factor *h*_*a*_, to account for factors such as Zelda, which do not exhibit spatial heterogeneity^88^. The variables *T*^*ab*^ and *E*^*αβ*^are coefficients that weight the transcriptional effect of *b* or *β* on *a* or *α*, respectively.

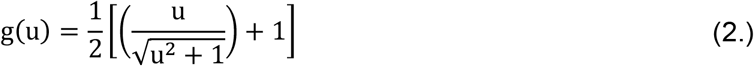

The second term simulates protein diffusion, such that *D*^*a*^ is the diffusion rate, and the third term stands for protein degradation, where *λ*^*a*^ is a measure of the half-life of *a*, such that 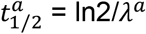. Finally, due to the additive property of enhancer-driven expression profiles, only the transcriptional weights are unique to each enhancer-driven profile, and the parameters for synthesis, diffusion, and degradation are unique to the protein (*b*) and not the species (*a*).

#### (2) Simulation Conditions

This model formulation consists of 68 variable parameters, of which 12 parameters modulate the synthesis, diffusion, and degradation rates of each of the four gap genes, and 56 parameters weigh the transcriptional effect of each species on another. For nc 13, ode15s integrates over the initial conditions (experimental expression profile for Hb and zeros for Kr, Gt, and Kni) for a simulation time of 16 mins from the start of nc 13, after which the embryo enters mitosis, and the synthesis rates are set to zero. After 21.1 mins from the start of nc 13, the embryo enters the interphase of nc 14, and the final conditions after the nc 13 mitosis are interpolated onto a new spatial grid of 100 points, which serves as the initial conditions of nc 14.

#### (3) Parameter Estimation

We implemented the Improved Stochastic Ranking Evolutionary Strategy *plus* (ISRES+) to estimate the 68 model parameters and fit the enhancer activity-informed model over the entire AP axis^51^. The data obtained from the Stark Lab datasets were interpolated onto a 71-point grid for nc 13 and a 100-point grid for nc 14. The model was then fit to these data using ISRES+, and the goodness of fit (GOF) was evaluated as the sum of the error normalized by the standard deviation during each iteration, such that a lower GOF represents a better fit. ISRES+ is a (μ-λ)-based evolutionary optimization algorithm that includes local search strategies such as linear regression and Newton’s method to efficiently converge to an optimum solution. The algorithm was run for 1000 generations with a population size of 350 and a recombination rate of 0.85. See Supplementary Section **5** for a description and value of the hyperparameters supplied to the model. This optimization strategy was repeated at least fifty times to estimate the 68 variable parameters in the model, of which ∼40 resulted in a GOF in the range 1-4.5, and of these six had a GOF less than 1.5.

#### (4) Simulating *Kr* mutants

To simulate *Kr* null mutants, we edited the EAGER model and removed all the equations and parameters associated with synthesis, diffusion, degradation, and interactions of *Kr*. The estimated parameters, generated by fitting the EAGER model to the gap protein expression data, were used to simulate gap protein expression profiles in a *Kr* null background. All the other conditions were kept the same as the EAGER simulations. Since the data available for gap protein expression in the *Kr* null background is for time class 3 (TC3) and time class 7 (TC7) of the nc 14, the model fits from the TC3 and TC7 are compared to their respective data. The results for identifiability and regularization studies also focus on TC3 and TC7, for the same reason.

### 6.3. Identifiability Analysis

#### (1) Regularization: LASSO & Ridge regression

These are regularized linear regression variants that attempt to minimize the GOF and reduce the number of variable parameters^76^. We incorporated these into the GOF evaluation function as an effort to reduce the number of transcriptional weights in the model.

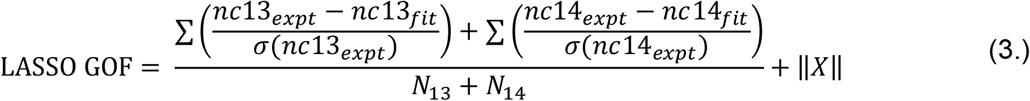

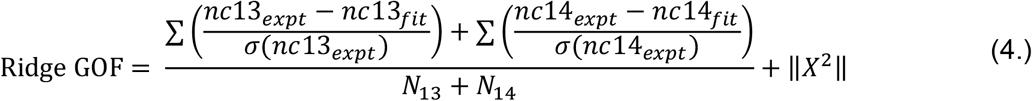

Here *N*_13_ and *N*_14_ denote the number of nuclei in nc 13 and 14 respectively, and *X* denotes all the transcriptional weights, including *T*^*a,b*^ and *E*^*αβ*^.

By reformulating the minimization problem (as Equation **3** and **4**) we implemented ISRES+ to generate a set of parameters which fit the model, see Methods: Parameter Estimation.

After regularization, an absolute cut-off of 0.05 was used to remove the transcriptional weights and simulate the reduced-EAGER model.

#### (2) Estimation of Confidence Intervals with “shuffled” parameter sets

The above parameter estimation and regularization strategies generated several parameter sets that adequately fit the experimental data and predicted the gap gene expression profiles in *Kr* null mutants. But the parameters have a wide distribution before and after implementing regularization strategies (Supplementary Section 1). To evaluate the robustness of these model predictions and the interdependencies among model parameters, we “shuffled” the parameters, as is a common practice in systems biology^89^. Using the six well-fit parameter sets with a GOF < 1.5, we generated 408 combinations. This was done by randomly “shuffling” each parameter while the other parameters in the 6 × (68 − 1) array were kept the same. The EAGER model was simulated at these 408 parameter sets, and the resulting mean (black curve) and confidence interval (±1 standard deviation; blue shaded region) were reported (Figure **2D-G;L-O**).

For the rEAGER model, 30 parameter sets with GOF in the range (1.6, 5.7) were “shuffled” one-by-one to generate 2040 parameter sets. The resulting mean (black curve) and confidence interval (green shaded region) were reported (Figure **4E-H;M-P**).

These 408 and 2040 parameter sets were used to simulate the predictions in the *Kr* null background for EAGER (Figure3D-F;J-L) and rEAGER (Figure **5D-F;J-L**), respectively.

#### (3) Sensitivity Analysis

Each of the 68 model parameters was increased by 1%, one at a time, to evaluate the effect of the individual parameter on peak characteristics of the gap gene expression profile, such as peak height (H), peak location (L), anterior peak boundary (A), and posterior peak boundary (P). This was done by adapting the findpeaks function in MATLAB to correctly identify each peak profile.

## 7. Acknowledgements

Portions of this research were conducted with the advanced computing resources provided by Texas A&M High Performance Research Computing.

## Notes

### Competing Interest Statement

The authors have declared no competing interest.

